# Importin-7-dependent nuclear localization of the *Flavivirus* core protein is required for infectious virus production

**DOI:** 10.1101/2023.10.11.561943

**Authors:** Yumi Itoh, Makoto Tokunaga, Yoichi Miyamoto, Tatsuya Suzuki, Akira Takada, Akinori Ninomiya, Mami Matsuda, Yoshihiro Yoneda, Masahiro Oka, Ryosuke Suzuki, Yoshiharu Matsuura, Toru Okamoto

## Abstract

*Flaviviridae* is a family of positive-stranded RNA viruses, including human pathogens such as Japanese encephalitis virus (JEV), dengue virus (DENV), zika virus (ZIKV), and West Nile virus (WNV). Nuclear localization of the viral core protein is conserved among *Flaviviridae*, and this feature may be targeted for developing broad-ranging anti-flavivirus drugs. However, the mechanism of core protein translocation to the nucleus and the importance of nuclear localization in the viral life cycle remain unknown. We aimed to identify the molecular mechanism underlying core protein nuclear translocation. We identified importin-7 (IPO7), an importin-β family protein, as a nuclear carrier for *Flaviviridae* core proteins. Nuclear import assays revealed that core protein was transported into the nucleus via IPO7, whereas *IPO7* deletion by CRISPR/Cas9 impaired their nuclear localization. To understand the importance of core protein nuclear localization, we evaluated the production of infectious virus or single-round-infectious-particles in wild-type or IPO7-deficient cells; both processes were significantly impaired in IPO7-deficient cells, whereas intracellular infectious virus levels were equivalent in wild-type and IPO7-deficient cells. These data suggested that IPO7-mediated nuclear localization of core proteins plays a role in the release of infectious virus particles of flaviviruses.

## INTRODUCTION

Viruses belonging to the family *Flaviviridae* including the genus *Flavivirus*, such as Japanese encephalitis virus (JEV), dengue virus (DENV), zika virus (ZIKV), and West Nile virus (WNV), are mosquito-borne human pathogens (Mukhopadhyay et al., 2005; Weaver et al., 2016). Infection with JEV or WNV can cause fatal neurological diseases in humans (Davis et al., 2006; Misra and Kalita, 2010). ZIKV is a neurotropic virus associated with Guillain–Barré syndrome, neuropathy, and myelitis in adults and children (Cao-Lormeau et al., 2016; Johansson et al., 2016). ZIKV infection during pregnancy can cause microcephaly in infants. DENV infection can cause dengue fever and dengue hemorrhagic fever (Halstead, 2007). The risk of infectious diseases caused by flaviviruses is increasing globally. DENV causes an estimated 390 million infections and 100 million symptomatic cases per year. Moreover, number of DENV infection and DENV-related death has been increasing in the last two decades (Pierson et al., 2020). Especially in Bangladesh, the highest number of dengue-related death was observed in 2022, compared to the previous years (Kayesh et al., 2023). It has been also reported that 70,000 cases of JEV infection have occurred in a year (Li et al., 2015). However, due to no cure for diseases caused by flavivirus infections, only relief of severe symptoms is conducted (Bifani et al., 2023). Therefore, prophylactic and therapeutic measures, including vaccines, are urgently needed.

*Flaviviridae* are single-stranded positive-sense RNA viruses. Viral RNA released from viral particles and is directly translated into a polyprotein of approximately 3,000 amino acids. The polyprotein is cleaved into 10 viral proteins by host or viral proteases. Viral replication and viral particle production occur in the endoplasmic reticulum (ER). Therefore, for many years, it was believed that the viral life cycle is completed in the cytoplasm. However, it is known that some flavivirus core proteins are translocated from the cytoplasm into the nucleus. Ala substitution in the JEV core protein at Gly42 and Pro43 impaired the nuclear localization of the core protein (Mori et al., 2005), and recombinant JEV possessing a core protein lacking the nuclear localization signal showed impaired propagation and attenuation in a mouse model. Mutations in the JEV core protein impaired the formation of viral infectious particles (Ishida et al., 2019). However, mutations of the core protein may affect the three-dimensional structure of the protein and have unknown side effects besides affecting nuclear localization. We recently showed that core protein nuclear localization is conserved among *Flaviviridae* and may be a target for broad-ranging anti-flavivirus drugs (Tokunaga et al., 2020); however, it remains unclear how the core protein is translocated from the cytoplasm to the nucleus.

Macromolecular transport between the cytoplasm and nucleus via the nuclear pore complex (NPC) is a fundamental process for information delivery for appropriate gene expression that is tightly regulated by importin/karyopherin family proteins (Wing et al., 2022). The importin/karyopherin family consists of more than 20 proteins that are categorized into two groups: importin-α and importin-β. Importin-α proteins are classical nuclear localization signal (NLS) receptors and function as adaptor molecules linking NLS-containing proteins with importin-β1, an importin-β family protein (Pumroy and Cingolani, 2015; Miyamoto et al., 2016). In the cytoplasm, NLS-containing proteins are recognized by importin-α and form a trimeric complex with importin-β1. The complex passes through the nuclear pore via interaction of importin-β1 with components of the NPC. In the nucleus, the GTP-bound form of Ran (RanGTP), a guanine nucleotide-binding protein that is abundant in the nucleus, strongly associates with importin-β1 to release the NLS-containing protein. Approximately 20 importin-β family members have been identified and function in nuclear import, export, or bidirectional transport (Chook and Suël, 2011; Kimura and Imamoto, 2014). For nuclear import, importin-β proteins bind directly to specific cargo proteins, and the complex passes through the NPC. Cargoes are released from importin-β proteins by binding with RanGTP in the nucleus.

In this study, we aimed to unravel the molecular mechanism underlying the nuclear localization of core proteins of flaviviruses. Specifically, we investigated the involvement of importin-7 (IPO7) in the nuclear transport of core proteins as well as the role of core protein nuclear localization in the flavivirus life cycle.

## RESULTS

### Importin-β-dependent nuclear localization of flavivirus core proteins

The core proteins of JEV, DENV, ZIKV, and WNV are localized not only in the cytoplasm, but also in the nucleolus (Mori et al., 2005; Tokunaga et al., 2020). In the present study, we used two different *in vitro* nuclear transport assays using recombinant core proteins. First, green fluorescent protein (GFP)-fused recombinant core proteins, with Alexa Fluor 555-conjugated antibodies as an injection marker, were injected into the cytoplasm of Huh7 cells using capillary needles. After the cells were incubated for 30 min, nuclear localization of the core proteins was observed based on GFP intensity (injection [IJ] assay). Second, digitonin-permeabilized Huh7 cells were incubated with GFP-fused core proteins and rabbit reticulocyte lysates and then, nuclear localization of the core proteins was observed by microscopy (permeabilization (PM) assay) (Figure 1A). The IJ assay revealed that GFP-fused core proteins of JEV, DENV, ZIKV, and WNV all translocated from the cytoplasm to the nucleus, whereas Ala substitution at Gly42 and Pro43 (GP) of the JEV core protein (GP/AA) abolished its nuclear localization (Figure 1B). In the PM assay, GST- and GFP-fused recombinant protein harboring a Simian virus-40 NLS sequence (SV40 NLS) and wild-type (WT) JEV core protein clearly localized in the nucleus, whereas the GP/AA JEV core mutant showed no nuclear localization (Figure 1C).

**Figure 1.**
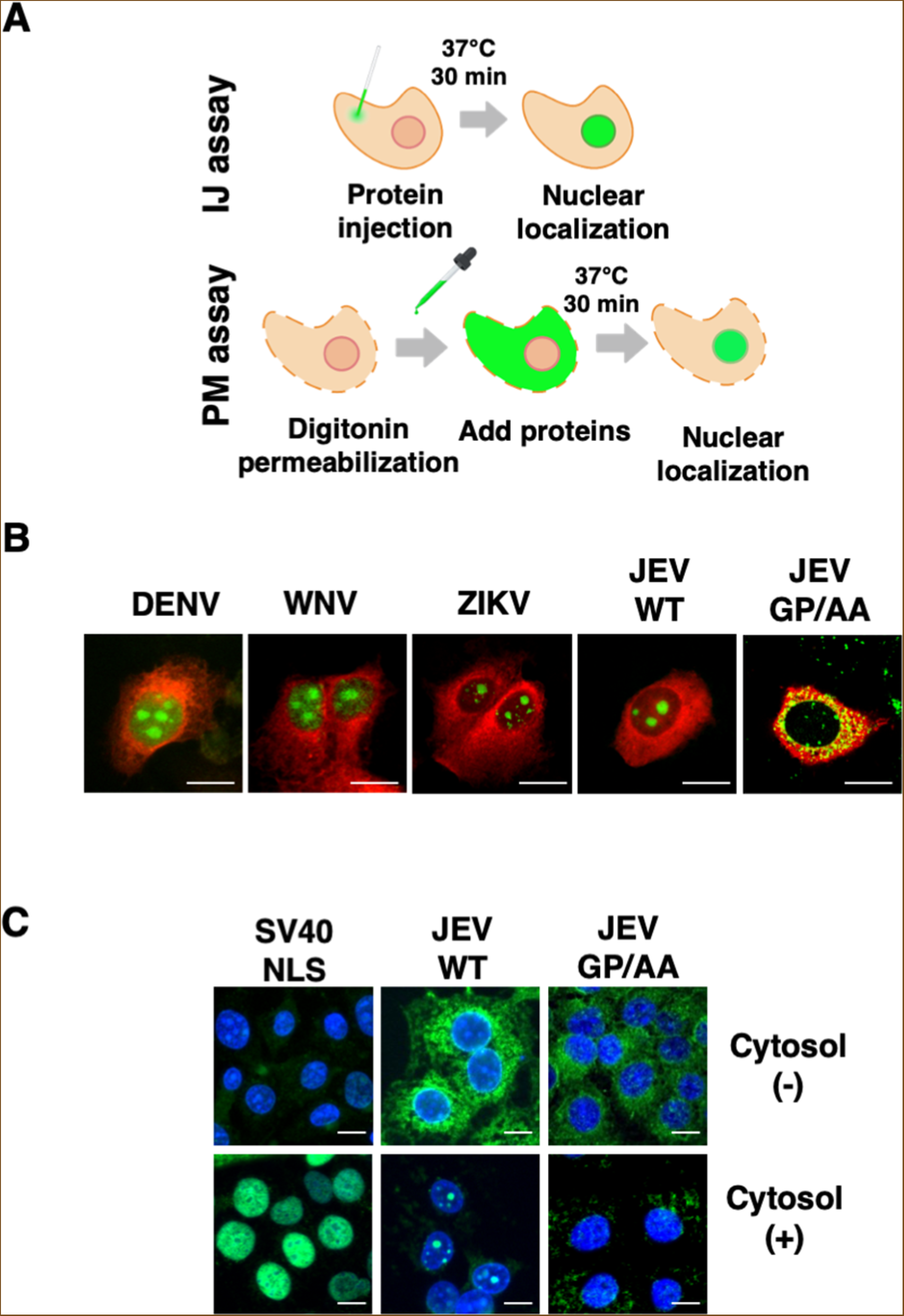
*In vitro* biochemical analysis of the nuclear localization of flavivirus core proteins. **(A)** Experimental design for analysis of the nuclear localization of core proteins. In the injection (IJ) assay, recombinant core proteins and immunoglobulin (IgG) were injected into Huh7 cells. After 30 min, the localization of recombinant core proteins was observed by confocal laser microscopy. In the permeabilization (PM) assay, Huh7 cells were treated with digitonin and the permeabilized cells were then incubated with recombinant core proteins. After 30 min, the localization of recombinant core proteins was observed by confocal laser microscopy. **(B)** Localization of core proteins of flaviviruses by the IJ assay. GFP-fused core protein and IgG labelled with Alexa Fluor 594 were injected into Huh7 cells. The scale bar indicates 20 μm. **(C)** PM assay using GST and GFP fused with the NLS of SV40 large T antigen (SV40 NLS) or WT or GP/AA core proteins of JEV. Rabbit reticulocyte lysate was employed as a cytosol source including import factors. The scale bar indicates 20 μm.

To understand the molecular pathway underlying nuclear transport of the core protein, we used several inhibitors. Bimax strongly binds to importin-α to inhibit importin-α-dependent nuclear localization (Kosugi, et al., 2008). Wheat germ agglutinin (WGA), an N-acetylglucosamine-binding lectin, inhibits active nuclear transport by binding to nucleoporins, which are heavily O-N-acetylglucosamine-modified (Li and Kohler, 2014). A dominant-negative Ran mutant mutated at Gln69 to Leu (RanGTP Q69L) and defective in GTP hydrolysis strongly binds to importin-β family proteins to inhibit importin-dependent nuclear localization (Dickmanns et al., 1996; Tachibana et al., 2000) (Figure 2A). We performed the IJ assay using the JEV core protein mixed with Bimax, Q69LRanGTP, or WGA. Injection with WGA inhibited core protein nuclear localization, indicating that it migrates into the nucleus via the NPC (Figure 2B). While Bimax showed no effect on core protein nuclear localization, RanGTP Q69L clearly retained the core protein in the cytoplasm (Figure 2B). This indicates that an importin-β-dependent pathway underlies the nuclear transport of the JEV core protein. As importin-β1 is a well-known nuclear transporter within the family, we first examined whether importin-β1 can directly bind to the core protein independent of importin-α. A glutathione S-transferase (GST) pull-down assay revealed that GST-importin-β1 did not interact with flag-tagged JEV core protein under the condition in which it bound to importin-α1 (Figure 2C). These data suggested that the JEV core protein is transported into the nucleus in an importin-β family-dependent manner, but not via importin-β1.

**Figure 2.**
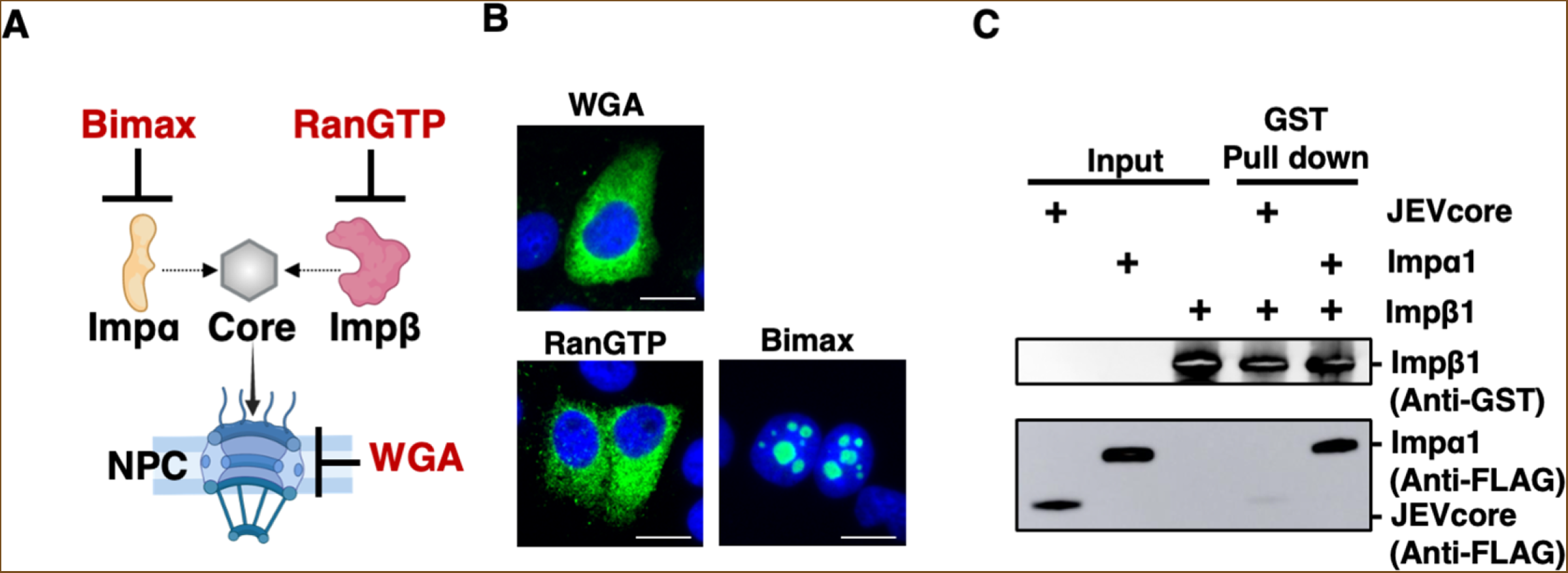
Nuclear pore complex (NPC)-mediated nuclear localization of core proteins is importin-α-independent and importin-β-dependent. **(A)** Scheme of the experiment used to assess the involvement of importin-α (impα), importin-β (impβ), and NPC. **(B)** The injection (IJ) assay was used to assess the involvement of importin-α, importin-β, and NPC in core protein nuclear localization. Bimax, RanGTP, and WGA were used as inhibitors of importin-α, importin-β, and nuclear pore complexes, respectively. The scale bar indicates 20 μm. **(C)** A GST-pull down assay was performed using GST-fused importin-β1 mixed with flag-tagged core protein or importin-α1, respectively. Input and immunoprecipitant samples were subjected to SDS-PAGE and detected by western blotting using anti-GST and anti-flag antibodies. The data are representative of three independent experiments.

### Identification of IPO7 as a carrier for core protein nuclear localization

We next attempted to identify proteins binding to the JEV core protein using mass spectrometry (MS). Strep tag-fused AcGFP or AcGFP-JEV core protein was expressed in 293T cells and immunoprecipitated using Strep-Tactin beads (Figure 3A). Among the core-binding proteins, we selected proteins belonging to the importin-β family, as shown in Figure 3B, and focused on IPO7 because it was identified the most peptides. To validate the interaction between IPO7 and the JEV core protein, we used an immunoprecipitation assay; IPO7 was pulled down with the AcGFP JEV core protein using an anti-flag antibody (Figure 3C).

**Figure 3.**
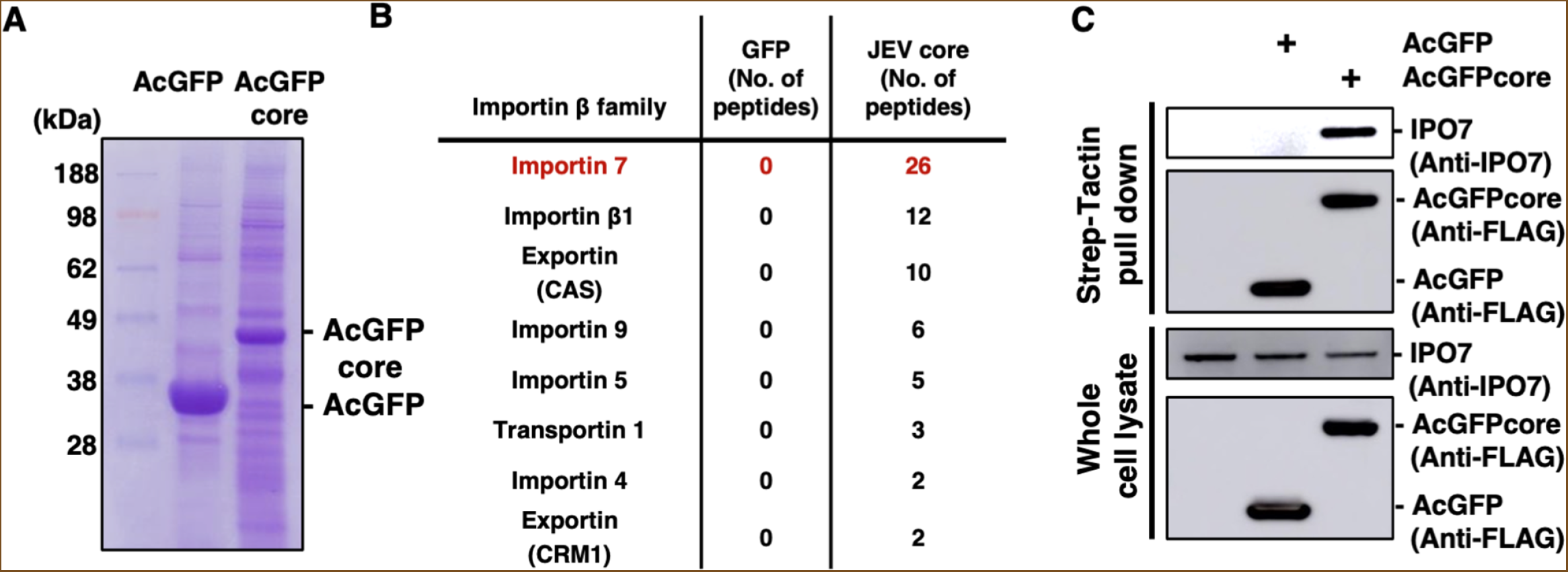
Importin-7 (IPO7) specifically interacts with the Japanese encephalitis virus (JEV) core protein. **(A)** GFP-FOS or AcGFP-JEV core-FOS was transfected into 293T cells and precipitated using Strep-Tactin Sepharose beads. The pull-down samples were analyzed by mass spectrometry (MS). **(B)** List of importin-β family proteins identified by MS analysis. The numbers of unique peptides identified by MS are indicated. **(C)** AcGFP-FOS or AcGFP-JEV core-FOS was transfected into 293T cells and precipitated using Strep-Tactin Sepharose beads. The precipitated samples were subjected to SDS-PAGE and detected by western blotting using anti-IPO7 and anti-flag antibodies.

To assess whether IPO7 binds directly to the core protein and then transports it into the nucleus, we purified bacterially expressed recombinant proteins and performed pull-down assays. As a positive control, we used ribosomal protein S7 (RPS7), which has been shown to be a specific cargo of IPO7, like RPL23A (Jäkel and Görlich, 1998). As shown in Figure 4A, AcGFP-RPS7 was pulled down with GST-IPO7 protein. The binding between IPO7 and RPS7 was disrupted upon the addition of RanGTP R69L. We also identified a direct interaction between IPO7 and the JEV core protein, and the addition of RanGTP clearly dissociated the binding even more effectively than for RPS7 (Figure 4B). These data suggested that the binding mode of IPO7 to the JEV core protein is similar to that to RPS7.

**Figure 4.**
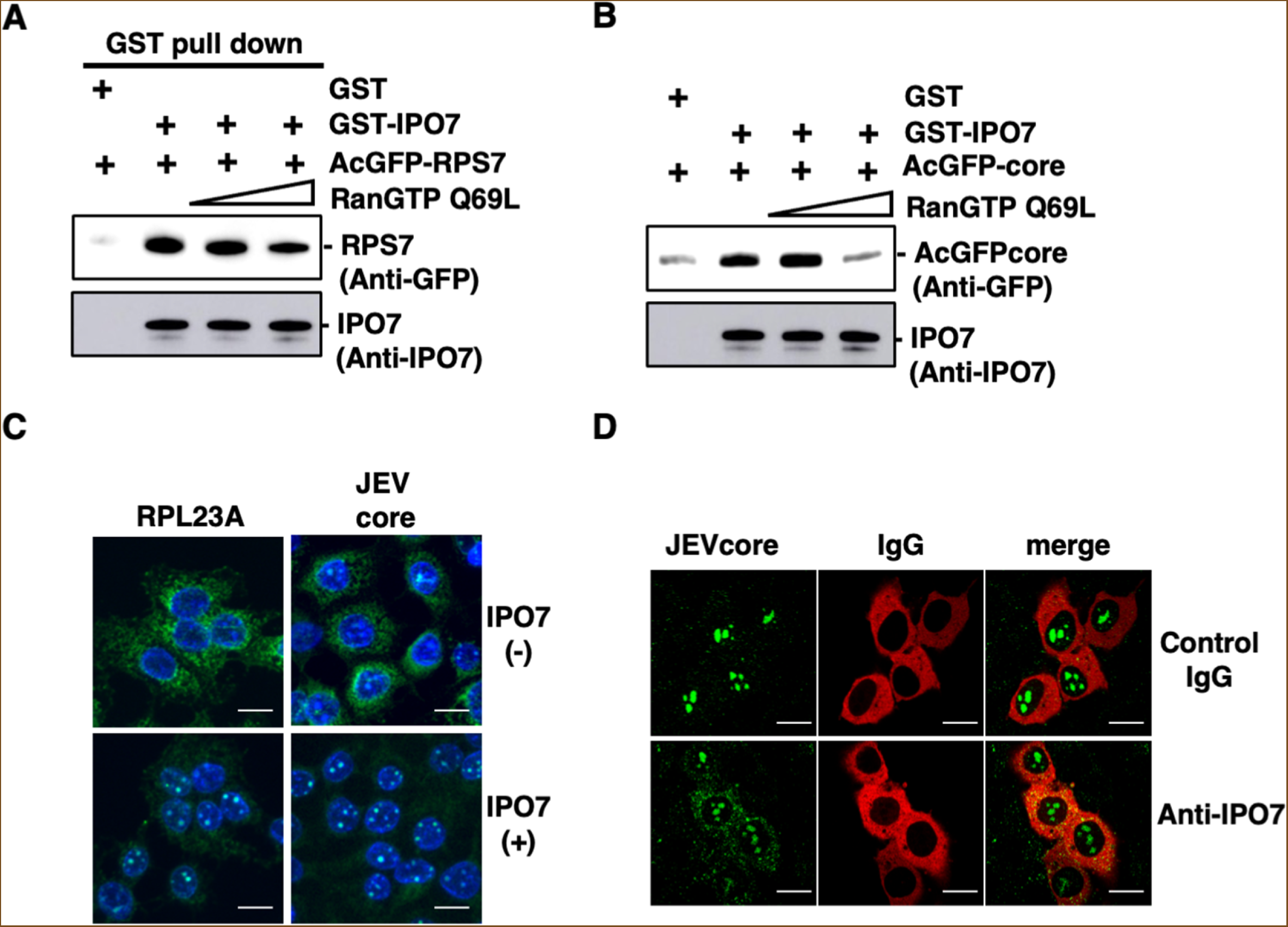
Importin-7 (IPO7) can transport flavivirus core proteins into the nucleus. **(A)** GST or GST-IPO7 was incubated with AcGFP-RPS7 in the presence or absence of 100 pmol or 770 pmol RanGTP Q69L. The pulled-down proteins were subjected to SDS-PAGE and detected by western blotting using anti-IPO7 and anti-GFP antibodies. **(B)** Either GST or GST-IPO7 was incubated with AcGFP-JEV core with or without 100 pmol or 770 pmol RanGTP Q69L. The pulled-down proteins were subjected to SDS-PAGE and detected by western blotting using anti-IPO7 and anti-GFP antibodies. **(C)** The permeabilization (PM) assay was performed using recombinant IPO7 protein. Digitonin-permeabilized Huh7 cells were incubated with GST-GFP-RPL23A or AcGFP-JEV core proteins with or without IPO7. Localization of core protein was observed by confocal laser microscopy. The scale bar indicates 20 μm. **(D)** IPO7 antibody or control IgG was injected with recombinant Japanese encephalitis virus (JEV) core protein and IgG conjugated with Alexa Fluor 594. Core protein localization was observed by confocal laser microscopy. The scale bar indicates 20 μm.

Finally, we examined whether IPO7 can transport the JEV core protein into the nucleus. A PM assay showed that the JEV core protein was localized in the nucleus when IPO7 recombinant protein was added, similar to the findings for RPL23A (Figure 4C). Microinjection of an antibody specific for IPO7 together with the JEV core protein into the cytoplasm significantly inhibited the nuclear localization of the JEV core protein (Figure 4D). These data showed that IPO7 acts as a nuclear carrier for the JEV core protein.

### IPO7 is a conserved nuclear carrier of flavivirus core proteins

Next, we generated IPO7-knockout (KO) cells using the CRISPR/Cas9 technology (Figure 5A). To examine the effect of IPO7 on the nuclear localization of core proteins of flaviviruses, recombinant core proteins of JEV, DENV, WNV, or ZIKV or SV40-NLS was injected into the cytoplasm of WT or IPO7-KO Huh7 cells. IJ assays showed that SV40-NLS was translocated into the nucleus in both WT and IPO7-KO Huh7 cells. The core proteins of JEV, DENV, and ZIKV were retained in the cytoplasm in IPO7-KO Huh7 cells, but not WT Huh7 cells (Figure 5B), while the core protein of WNV was partially retained in the cytoplasm in IPO7-KO Huh7 cells. Consistent with this finding, AcGFP-core protein plasmid transfection showed that nuclear localization of the JEV, DENV, WNV, and ZIKV core proteins was partially inhibited in IPO7-KO Huh7 cells but not in WT Huh7 cells (Figure 5C). We also examined core protein localization in JEV-infected IPO7-KO Huh7 cells. Nuclear localization of core protein was observed WT Huh7 cells but not in IPO7-KO Huh7 cells (Figure 5D). Although viral titers were lower in IPO7-KO Huh7 cells compared with those in WT Huh7 cells, intracellular viral RNA levels in IPO7-KO Huh7 cells were significantly higher than those in WT Huh7 cells (Figure 5E and 5F). We next quantified intracellular infectious virus particles upon repeated freezing and thawing of virus-infected cells. We found that intracellular infectious virus levels were equivalent in IPO7-KO and WT Huh7 cells (Figure 5G). Finally, we examined the effect of IPO7 on the life cycles of other flaviviruses, including DENV and ZIKV. Infectious viral titers were significantly decreased in supernatants of ZIKV- and DENV-infected IPO7-KO Huh7 cells compared to those of infected WT cells (Figure 5H). Next, we established additional IPO7-KO Huh7 cell lines and IPO7-KO 293T cells to analyze the effect of IPO7 KO in other cell types. Three additional IPO7-KO Huh7 cell lines also showed impaired core protein nuclear localization and viral replication (Figure 6A–C). The IPO7-KO 293T cells showed significantly impaired JEV propagation (Figure 6D). These data suggested that IPO7 is a conserved nuclear importer of core proteins of *Flaviviridae* and plays an important role in the life cycle of these viruses.

**Figure 5.**
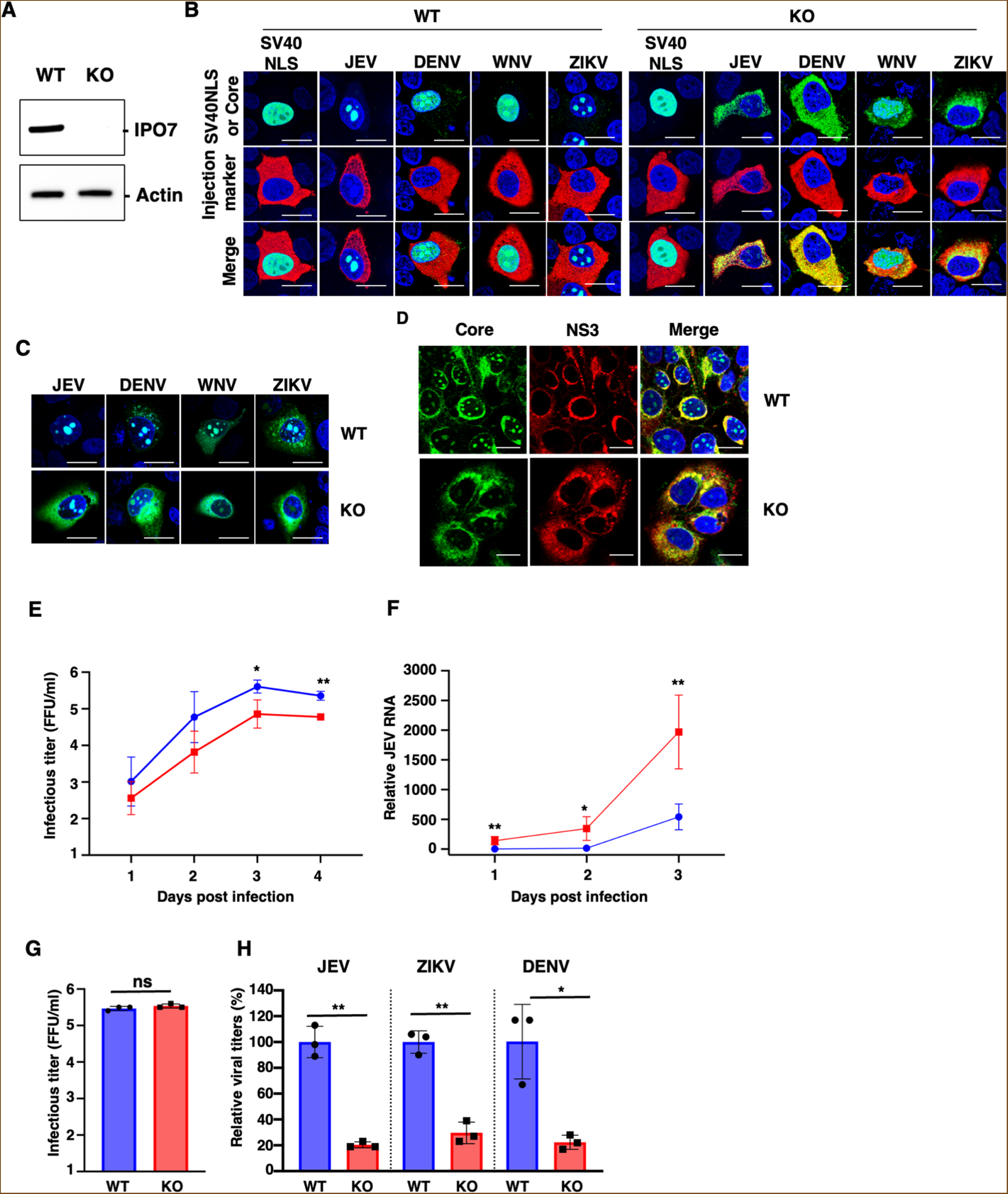
Importin-7 (IPO7) deficiency in Huh7 cells impairs the nuclear localization of core proteins of *Flaviviridae*. **(A)** IPO7 expression in IPO7-KO Huh7 cells was confirmed by western blotting. **(B)** GST and GFP fused the SV40 large T antigen NLS or GFP fused core proteins of Flaviviridae were microinjected into the the cytoplasm of WT or KO cells with red fluorescence protein-conjugated antibody as a cytoplasmic injection marker. The scale bar indicates 20 μm. **(C)** Plasmids encoding GFP fused with core proteins of *Flaviviridae* were transfected into wild-type (WT) or IPO7-KO Huh7 cells. The scale bar indicates 20 μm. **(D)** The WT or IPO7-KO Huh7 cells were infected with Japanese encephalitis virus (JEV). Subcellular localization of Core protein or NS3 were observed using antibodies against for JEV core or NS3, respectively. The scale bar indicates 20 μm. **(E–G)** WT or IPO7-KO Huh7 cells were infected with JEV. The viral titers in supernatants (E), intracellular viral RNA (F), and intracellular viral titers (G) were quantified. **(H)** WT or IPO7-KO Huh7 cells were infected with JEV, zika virus (ZIKV), or dengue virus (DENV). At 2 dpi, viral titers in supernatants were determined by the focus-forming unit (FFU) assay. Data are representative of two (A-D) independent experiments and are presented as the mean ± SD (E–H). Significance (\**p* < 0.05; \*\**p* < 0.01; n.s., not significant) was determined using Student’s *t*-test (*n* = 3).

**Figure 6.**
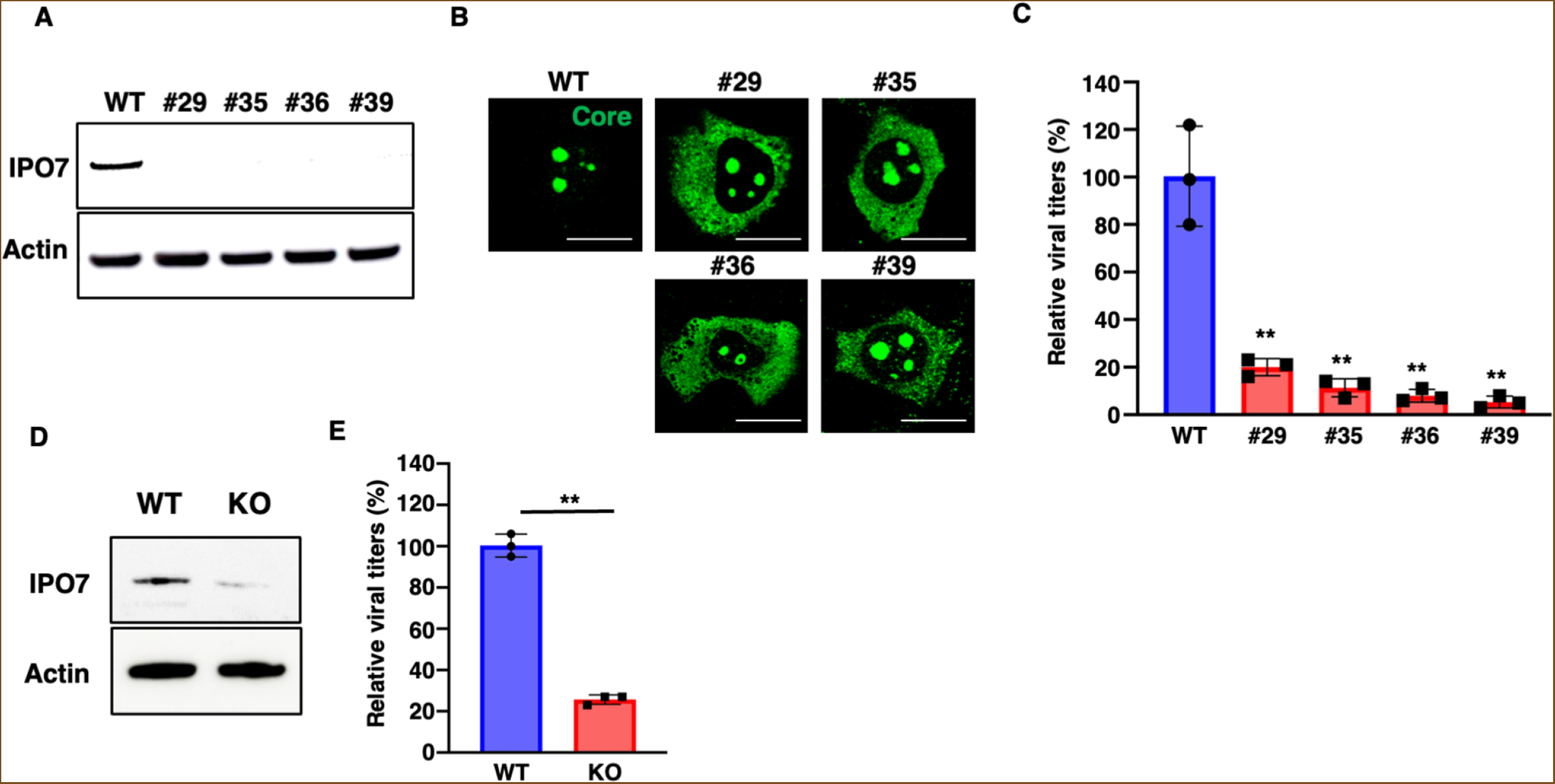
Four importin-7 knockout (IPO7-KO) Huh7 clones showed impaired nuclear localization of the core protein and viral propagation. **(A)** IPO7 expression in IPO7-KO Huh7 cell clones was confirmed by western blotting. **(B)** The injection (IJ) assay was performed using GFP fused with core proteins of *Flaviviridae* in wild-type (WT) or IPO7-KO Huh7 cells. The scale bar indicates 20 μm. **(C)** WT or IPO7-KO Huh7 cells were infected with Japanese encephalitis virus (JEV). At 2 dpi, viral titers in supernatants were determined by the focus-forming unit (FFU) assay. **(D)** IPO7 expression in IPO7-KO 293T cells was confirmed by western blotting. **(E)** WT or IPO7-KO 293T cells were infected with JEV. At 2 dpi, viral titers in supernatants were determined by the FFU assay. Data are representative of two (A, B) independent experiments and are presented as the mean ± SD of three independent experiments (C). Significance (\**p* < 0.05; \*\**p* < 0.01; n.s., not significant) was determined using Student’s *t*-test (*n* = 3).

### IPO7 is involved in the release of infectious virus particles of *Flaviviridae*

The above data suggested that IPO7-dependent nuclear localization of flavivirus core proteins may play a role in virus release. To examine the importance of nuclear localization in the viral life cycle, we employed single-round infectious particles (SRIPs) of flaviviruses (Matsuda et al., 2018, Yamanaka et al., 2021). Three kinds of plasmids, Replicon, core, and prME expression plasmids were transfected into packaging cells to produce SRIPs. The SRIPs obtained were used to infect Vero cells, and NanoLuc luciferase (Nluc) values of the SRIP-infected Vero cells were quantified (Figure 7A). SRIPs were generated using a yellow fever virus (YFV)-derived replicon and prME plasmid, and WT JEV core or GP/AA JEV core plasmid. Nluc values in Vero cells infected with GP/AA JEV core SRIPs were significantly lower than those in cells infected with WT JEV core SRIPs (Figure 7B). This suggested that nuclear localization of core proteins was involved in the release of infectious virus particles. Next, we examined the involvement of IPO7 in viral RNA replication. *In vitro*-transcribed RNA was electroporated into IPO7-KO or WT Huh7 cells, and Nluc values were monitored. Consistent with the data in Figure 5F, Nluc values in IPO7-KO Huh7 cells were significantly higher than those in WT Huh7 cells (Figure 7C). We then evaluated SRIP production in the IPO7-KO and WT Huh7 cells. Nluc values in Vero cells infected with JEV SRIPs of IPO7-KO Huh7 cells were significantly lower than those in Vero cells infected with JEV SRIPs of WT Huh7 cells (Figure 7D). Similar results were obtained when using DENV, ZIKV, and YFV SRIPs (Figure 7E). Finally, we confirmed that the numbers of intracellular infectious viruses were equivalent between IPO7-KO or WT Huh7 cells (Figure 7F). Taken together, the data suggested that IPO7-mediated core protein nuclear localization is important for the release of infectious flavivirus particles.

**Figure 7.**
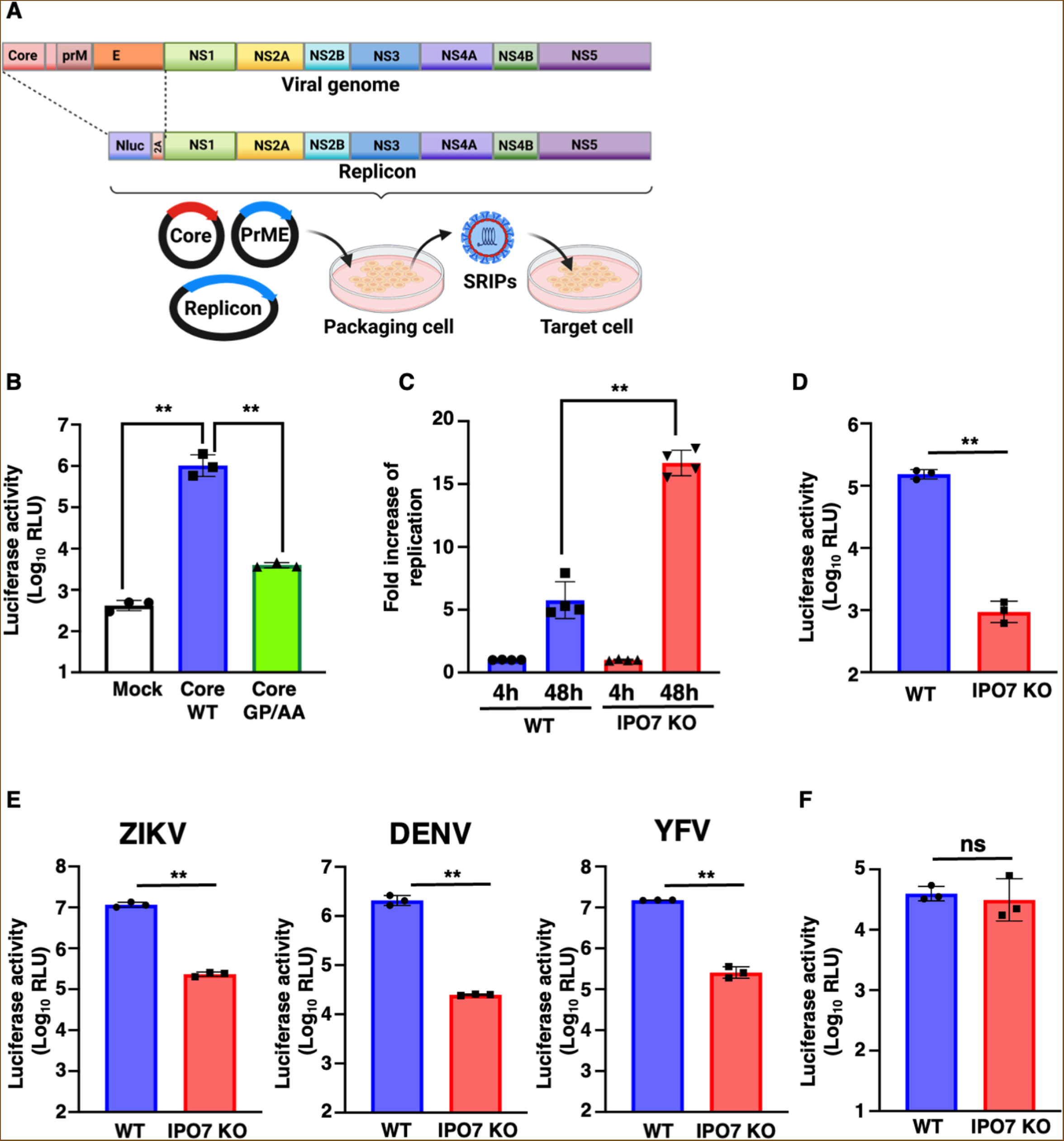
Disruption of the nuclear localization of the core proteins of *Flaviviridae* impairs efficient infectious virus release. **(A)** Schematic procedure of the single-round infectious particle (SRIP) generation experiments. **(B)** Three different plasmids containing the core, prME, and replicon were transfected into 293T cells to produce SRIPs. SRIPs produced by wild-type (WT) or GP/AA core protein were used to infect Vero cells. Nluc reporter activity was quantified using the Nano-Glo Luciferase Assay System. **(C)** *In vitro*-transcribed Japanese encephalitis virus (JEV) replicon RNA was electroporated into WT and IPO7-KO Huh7 cells. Viral RNA replication capacity is shown as relative reporter activity normalized to reporter activity at 4 h post electroporation. **(D)** JEV SRIPs were generated from WT or IPO7-KO Huh7 cells. The reporter activity of the SRIPs was determined after infection into Vero cells. **(E)** SRIPs of zika virus (ZIKV), dengue virus (DENV), or yellow fever virus (YFV) were generated from WT or IPO7-KO Huh7 cells. The reporter activity of the SRIPs was determined after infection into Vero cells. **(F)** JEV SRIPs were generated from WT or IPO7-KO Huh7 cells. Intracellular SRIPs were collected after each of repeated freeze–thaw cycles. The reporter activity of the SRIPs was determined after infection into Vero cells. Data are presented as the mean ± SD of three independent experiments. Significance (\*\**p* < 0.01; n.s., not significant) was determined using Student’s *t*-test (*n* = 3; B, D–F, n = 4; C).

## DISCUSSION

This study aimed to elucidate the molecular mechanism underlying the nuclear translocation of flavivirus core proteins and the relevance of this translocation to the viral life cycle. We identified IPO7, also known as RanBP7, as a nuclear transporter of core proteins of flaviviruses such as DENV, ZIKA, WNV, and JEV. Although nuclear transport of WNV core protein was partially inhibited in IPO7-KO Huh7 cells, our data showed IPO7 is involved in a nuclear transport of core proteins of *Flaviviridae* family. Because it has been reported that interaction between WNV core protein and importin-α/β enables WNV to produce viral particles efficiently (Bhuvanakantham et al., 2009), nuclear localization of WNV core protein was regulated by several nuclear carriers. IPO7 is an importin-β family member and can transport specific cargos, including ribosomal proteins such as L23a, S7, and L5 (Jäkel and Görlich, 1998), histones (Baake et al., 2001; Mühlhäusser et al., 2001), thyroid hormone receptor α1 (TRα1) (Roggero et al., 2016), Smads (Xu et al., 2007; Yao et al., 2008), Erk (Lorenzen et al., 2001), and YAP (García et al., 2022). IPO7 has also been shown to transport histone H1 in a heterodimer with importin β1 into the nucleus (Jäkel et al., 1999; Bäuerle et al., 2002). Regarding viral infection and replication, IPO7 has been reported to regulate viral proteins, such as HIV-1 integrase (Ao et al., 2007) or purified intracellular reverse transcription complex (Fassati et al., 2003), and inhibit virus replication. Using stable IPO7-knockdown cells, the nuclear entry of viral genomic DNA has been demonstrated to involve IPO7 (Zaitseva et al., 2009), indicating the physiological significance of IPO7 in the life cycle of HIV-1. We found here that IPO7-KO Huh7 cells showed impaired infectious virus production of flaviviruses. Similar results were obtained with GP/AA core protein (Figure 7B) and in a previous study (Ishida et al., 2019), suggesting that core protein nuclear localization plays a role in viral release. In contrast, viral RNA replication was increased in IPO7-KO Huh7 cells. Whether IPO7 may transport other factors possessing antiviral activity against flaviviruses remains unclear and requires further studies.

There is growing evidence that the consensus NLS sequence recognized by IPO7 is enriched in basic amino acids (EKRKI(E/R)/(K/L/R/S/T)) (Panagiotopoulos et al., 2021). Although no corresponding sequences are found in flavivirus core proteins, they harbor characteristic basic amino acid sequences, particularly in the C-terminus (Poonsiri et al., 2019). Indeed, the NLS sequences of WNV is localized in the amino acid region 85–101 and is recognized by importin-α (Bhuvanakantham et al., 2009). In DENV, three NLSs, ^6^KKAR^9, 73^KKSK^76^, and a bipartite NLS ^85^RKeigrmlnilnRRRR^100^, have been reported (Bulich and Aaskov 1992; Wang et al., 2002; Sangiambut et al., 2008). In contrast, in our study, IJ experiments showed no effect of Bimax, an importin-α inhibitor, on the nuclear entry of the JEV core protein (Kosugi et al., 2008). Considering the cytoplasmic localization of all flavivirus core proteins assessed in IPO7-KO cells, we suggest that IPO7 serves as a common nuclear transporter for core proteins of flaviviruses.

While RNA viruses, including flaviviruses, are considered to replicate in the cytoplasm of mammalian cells, increasing evidence suggests nucleolar localization of the viruses (Rawlinson and Moseley, 2015). It is highly interesting to know the relationship between the nucleolar localization of core proteins and their function in virus replication. Among the 20 importin-β family members, flaviviruses preferentially use IPO7 as a core protein nuclear transporter. There may be a physiological association in that IPO7 may transport several ribosomal proteins into the nucleus and may thus be involved in ribosomal biogenesis (Jäkel and Görlich, 1998; Golomb et al., 2012). In addition, IPO7 has been shown to inefficiently release the ribosomal protein RPL23a, even in the presence of RanGTP, suggesting the presence of other molecules that can release the IPO7-ribosomal protein complex (Jäkel and Görlich, 1998). In our binding assay, the IPO7–RPS7 bond dissociated inefficiently despite the addition of excess amounts of RanGTP Q69L. It may be a candidate associating protein of nucleolar phosphoprotein B23, which we previously identified to bind with the JEV core protein (Tsuda et al., 2006). Although it is unclear whether B23 is a target of core protein localizing in the nucleolus, interactome analysis may clarify why flavivirus core proteins are specifically recognized by IPO7 and transported into the nucleolus.

Alternatively, the nucleolar structure has also been considered to guide the nucleolar targeting of core proteins. We previously reported that several compounds, such as CDK inhibitors, dramatically disrupted the nucleolar morphology and significantly suppressed viral replication (Tokunaga et al., 2020). These and our current data suggest that a functional correlation between the nucleolar structure and core protein might be required for proper capsid formation or envelopment and resulted in production of infectious virus. The nucleolus is known to be generated by phase separation (Latonen 2019). Further, flavivirus capsid proteins show phase separation characteristics (Ambroggio et al., 2021; Saito et al., 2021). We assume that core protein nucleolar localization may occur via phase separation in the nucleolus. Further studies are required to better understand the physiological functions of flavivirus core proteins in the nucleus and nucleolus.

In summary, it is the first report identify that nuclear localization of core protein is mediated by IPO7 and plays an important role in the propagation of flaviviruses. These findings have contributed to further understand the flavivirus life cycle and its possibility as therapeutic target. However, the function of core protein in the nucleus and nucleolus is still unclear. Further work will be needed to unveil more details of physiological function of core proteins in the nucleus that lead to the efficient release of infectious viral particles. This approach might identify a novel drug target for flavivirus infectious diseases. Therefore, it is needed to identify the condition to inhibit the nuclear localization of core proteins and evaluate its importance in vivo.

## MATERIALS AND METHODS

### Viruses and cells

293T (ATCC), Huh7 (JCRB0403, JCRB Cell Bank), and Vero (JCRB0111) cells were cultured in Dulbecco’s modified Eagle’s medium (DMEM, Wako) containing 10% fetal bovine serum (FBS) (Gibco) and penicillin/streptomycin (100 U/mL, Invitrogen) (P/S) at 37 °C in the presence of 5% CO_2_. C6/36 (IFO50010), a Singh’s *Aedes albopictus* cell clone, were maintained in Leibovitz’s L-15 medium (Thermo Fisher Scientific) supplemented with 10% FBS, 2.4 g/L tryptose phosphate broth (BD), and P/S. *Spodoptera frugiperda* (Sf-9) cells were maintained in Sf-900 II SFM medium (Thermo Fisher Scientific). All cell lines were routinely tested to be negative for mycoplasma contamination. JEV (strain AT31), DENV (strain 16688), and ZIKV (strain MR766) were obtained from National Institute of Infectious Diseases (NIID) and maintained in C6/36 cells. Infectious titers of JEV, DENV, and ZIKV were determined using the focus-forming unit (FFU) assay.

### Plasmid generation

cDNAs of the core proteins of JEV (strain AT31), DENV (strain 16681), and ZIKV (strain MR766) were PCR-amplified from cDNA from virus-infected cells, and cDNA of the WNV (strain NY99) core protein was PCR-amplified from cDNA synthesized at IDT DNA Technologies (Coralville, IA, USA). The cDNAs were cloned into pCAGGS or pFastBac-1 together with AcGFP and a flag-<u>o</u>ne-<u>s</u>trep (FOS) tag. *IPO7* cDNA was PCR-amplified using primers 5′-GACCCAAGGAGGATCCATGGACCCCAACACC-3′, 5′-TCTAGAGTCGCGGCCGCTCAATTCATCCCTGG-3′ and cloned into pGEX6P-2 (Cytiva, WA, USA). The relevant oligonucleotide of SV40 large T antigen NLS (PPKKKRKVED) was ligated into the pGEX2T vector (GE Healthcare) carrying the *GFP* gene at the C-terminus of the multi-cloning site to produce GST-SV40NLS-GFP protein (referred to as SV40 NLS). Plasmids for GST-fused and flag-tagged importin-α1 (KPNA2), importin-β1, WT Ran, and Q69LRan were obtained as described previously (Sekimoto et al., 1997; Miyamoto et al., 2002; Miyamoto et al., 2013; Kimoto et al., 2015). RPL23A and RPS7 cDNAs were amplified from cDNA of Huh7 cells and cloned into pGEX6P-2. sgRNAs of human IPO7 (5′-ATGATCGACCTGAGTTACCA-3′), (5′-AATACATACCTGATGAGCTC-3′) and (5′-CCTGGTGCTGTTTCTCGATC-3′) were cloned into pX330. The plasmids for SRIPs have been reported previously (Matsuda et al., 2018). All cloning experiments were performed using an In-Fusion HD Cloning Kit (Takara Bio Inc., Shiga, Japan), and plasmid sequences were confirmed by DNA sequencing (at the Core Instrumentation Facility, Research Institute for Microbial Diseases, Osaka University, Osaka, Japan).

### Purification of recombinant proteins

Bacterially expressed recombinant proteins fused to GST were purified as described previously (Miyamoto and OKa, 2016). Importin-α and Ran proteins lacking GST were prepared by cleavage with PreScission protease (20 U/mg of fusion protein; GE Healthcare, Uppsala, Sweden) at 4 °C for 12 h. The GST-cleaved Ran protein was equilibrated with buffer (50 mM HEPES [pH 7.3], 75 mM NaCl, 1 mM MgCl_2_, 1 mM dithiothreitol [DTT], and 0.1 mM phenylmethylsulfonyl fluoride) and incubated with GDP or GTP (2 mM; Sigma-Aldrich, St. Louis, MO, USA) and WT RanGDP or Q69LRanGTP on ice for 1 h. To stop the reaction, the proteins were incubated with 50 mM MgCl_2_. The proteins were equilibrated with dialysis buffer (20 mM HEPES [pH 7.3]), 100 mM CH_3_COOK, 2 mM DTT, and protease inhibitor (Protease Inhibitor Cocktail, Nacalai Tesque, Kyoto, Japan) using an Amicon Ultra device (10K; Millipore, Billerica, MA, USA).

### Generation of recombinant core proteins using an insect expression system

Recombinant baculoviruses (AcNPV) were generated using pFastBac1 AcGFP-JEVcore-FOS and a Bac-to-Bac system (Invitrogen, Carlsbad, CA, USA) following the manufacturer’s protocol. Recombinant baculoviruses carrying JEV core protein (JEVcore) cDNA were propagated in Sf-9 cells. To generate recombinant JEV core protein, Sf-9 cells (2 × 10^6^) were seeded in 10-cm dishes (Greiner Bio-One GmbH, Frickenhausen, Germany) and were infected with recombinant AcNPV at a multiplicity of infection of 10 and incubated for 3 days. Cells were collected and lysed in lysis buffer (20 mM Tris-HCl [pH 7.4], 135 mM NaCl, 1% Triton X-100, 1% glycerol, and protease inhibitor cocktail tablets [Roche Diagnostics, Mannheim, Germany]) at 4°C for 30 min, followed by centrifugation at 14,000 × *g* at 4°C for 15 min. The supernatants were incubated with Strep-Tactin Sepharose (IBA GMBH) overnight. The beads were washed using Strep-tag washing buffer (IBA GMBH) three times and eluted using elution buffer (IBA GMBH) containing D-desthiobiotin. The recombinant proteins were concentrated using an Amicon Ultra (10K; Millipore, Billerica, MA, USA).

### Microinjection

Huh7 cells were seeded on coverslips (MATSUNAMI) in 35-mm dishes (Corning, Acton, MA, USA). Within 1 min, AcGFP-JEVcore recombinant protein and Alexa Fluor 594 or 555-conjugated antibody (2 mg/mL, Thermo Fisher Scientific, as an injection marker) were microinjected into the cytoplasm at a final concentration of 1 mg/mL. After incubation for 30 min, the cells were fixed with 3.7% formaldehyde in PBS.

### Semi-intact nuclear import assay

Huh7 cells were treated with 40 μg/mL digitonin (Nacalai Tesque) on ice for 5 min and washed twice with the transport buffer (20 mM HEPES [pH 7.3], 110 mM CH_3_COOK, 5 mM CH_3_COONa, 2 mM (CH_3_COO)_2_Mg, 0.5 mM EGTA, 2 mM DTT, and protease inhibitor [Nacalai Tesque]) to minimize residual protein content in the cytoplasm. The permeabilized cells were incubated in ice-cold transport buffer for 10 min and then incubated at 37 °C for 30 min in the following transport mixtures: for the experiment shown in Figure 1C, 4 pmol of SV40 NLS protein, 20 pmol of AcGFP-JEVCoreWT, AcGFP-JEVCoreGP/AA, or 10 pmol of GST-GFP-RPL23A, Rabbit Reticulocyte Lysate (Promega), ATP regeneration system (0.5 mM ATP [Wako]), 20 U/mL creatine phosphokinase (Sigma-Aldrich), and 5 mM creatine phosphate (Sigma-Aldrich); for the experiment shown in Figure 4C, 4 pmol of IPO7 and 40 pmol of WT RanGDP with ATP regeneration system in a total volume of 10 μL per sample. After incubation, the cells were fixed with 3.7% formaldehyde in PBS. The cells were observed under a Nikon Eclipse 80i microscope (Nikon, Tokyo, Japan).

### Immunofluorescence

Cells were fixed with 3.7% formaldehyde in PBS at room temperature for 15 min. After washing in PBS, the cells were treated with 0.1% Triton X-100 in PBS for 5 min, blocked in PBS containing 5% skim milk for 30 min, and incubated in appropriately diluted (in PBS containing 5% skim milk) primary antibodies at room temperature for 2 h or at 4 °C overnight. Then, the cells were incubated with Alexa Fluor-conjugated secondary antibodies (fluorescence at 488, 594, 555, or 645 nm; Thermo Fisher Scientific, Rockford, IL, USA). Nuclei were counterstained with DAPI (1:10,000 in PBS; Dojindo Laboratories, Kumamoto, Japan) for 10 min. The samples were examined using a Zeiss Axiophot fluorescence microscope (Carl Zeiss, Gottingen, Germany) and a Nikon Eclipse 80i microscope (Nikon).

### Antibodies

The following antibodies were used in this study: mouse monoclonal antibodies against JEV/DENV/ZIKV NS3 (clone 578) (Suzuki et al., 2018), anti-IPO7 (ab99273; Abcam, Cambridge, UK), anti-JEV core antibody (GTX131368; GeneTex, Irvine, CA, USA), and anti-flag monoclonal antibody (clone M2; Sigma, St. Louis, MO, USA).

### Virus infection

Huh7 cells were seeded in 24-well plates at 3 × 10^4^ cells/well and incubated for 24 h. Then, the cells were infected with JEV, DENV, or ZIKV at a multiplicity of infection of 5 for 2 h. The virus-infected Huh7 cells were washed with PBS and cultured in culture medium. Viral infectivity was determined as FFUs using an immunostaining assay.

### FFU assay

The FFU assay was performed as described previously (8). Culture media from Huh7 cells infected with JEV were collected and serially diluted. The diluted media including viruses were added to Vero cells seeded in 24-well plates (5 × 10^4^ cells/well). After 2 h of adsorption, the cells were washed with PBS followed by culture in DMEM containing 10% FBS, 100 U/mL penicillin and 100 μg/mL streptomycin, and 1.25% methylcellulose 4,000cP (M0512, Merck KGaA, Darmstadt, Germany). Two days later, the cells were fixed with 4% paraformaldehyde and permeabilized with 0.5% Triton X-100 for 5 min. The cells were incubated with an anti-NS3 (clone 578) antibody (54) for 30 min and then with a biotin-conjugated anti-mouse IgG secondary antibody (Vector Laboratories, Burlingame, CA, USA) for 30 min. After washing with PBS, infectious foci were visualized by incubation with Streptavidin Biotin Complex Peroxidase (Nacalai Tesque) for 30 min followed by incubation with VIP Peroxidase Substrate (Vector Laboratories) for 10 min.

### Western blotting

Cell lysates were prepared by incubation in Onyx lysis buffer (20 mM Tris-HCl [pH 7.4], 135 mM NaCl, 1% Triton X-100, 1% glycerol, and protease inhibitor cocktail tablets [Roche Molecular Biochemicals]) at 4 °C for 15 min. After sonication, the lysates were centrifuged at 13,000 x g at 4 °C for 5 min. Protein concentrations were determined using Protein Assay Dye Reagent Concentrate (Bio-Rad). Supernatants were incubated with sample buffer at 95°C for 5 min. The samples (50 μg of proteins) were resolved by SDS-PAGE (Novex/Life Technologies) and transferred onto nitrocellulose membranes (iBlot/Life Technologies). The membranes were blocked with PBS containing 5% skim milk and incubated with primary antibodies at 4 °C overnight. After washing, the membranes were reacted with HRP-conjugated secondary antibody at room temperature for 2 h. The immune complexes were visualized using Super Signal West Femto substrate (Pierce) and detected using an Amersham ImageQuant 800 system (Cytiva Life Science, Marlborough, MA, USA).

### MS

293T cells transfected with pCAG AcGFP-FOS or pCAG AcGFP-JEVcore-FOS were incubated for 48 h and lysed in lysis buffer (20 mM Tris-HCl [pH = 7.4], 135 mM NaCl, 1% Triton X-100, 1% glycerol, and protease inhibitor cocktail tablets [Roche]). The cell lysates were incubated with Strep-Tactin Sepharose (IBM GmbH) for 90 min and washed three times with lysis buffer. The precipitates were boiled in sample buffer at 95 °C for 5 min as described previously. The proteins were subject to SDS-PAGE, stained with Coomassie brilliant blue, purified, and divided into 10 aliquots. The proteins were reduced with 10 mM DTT, alkylated with 55 mM iodoacetamide, and digested with trypsin (Promega). The peptides were subjected to MS using an LC-ESI system. MS data were obtained by nanocapillary reversed-phase LC-MS/MS using a C18 column (0.1 × 150 mm) on a nanoLC system (Advance/Michrom BioResources) coupled to an LTQ Orbitrap Velos mass spectrometer (Thermo Fisher Scientific). The mobile phase consisted of water containing 0.1% formic acid (solvent A) and acetonitrile (solvent B). Peptides were eluted using a gradient of 5–35% B for 45 min at a flow rate of 500 nL/min. The mass scanning range of the instrument was set to m/z 350–1,500. The ion spray voltage was set to 1.8 kV in the positive ion mode. The MS/MS spectra were acquired by automatic switching between MS and MS/MS modes (with collisional energy set to 35%). The dynamic exclusion was set to 30 s, with 10 ppm tolerance. Helium gas was used as a collision gas. The data obtained were processed using Proteome Discoverer (Thermo Fisher Scientific) and peptides were identified using MASCOT (Matrix Science, UK) against the Swiss-Prot human database. Precursor mass tolerance was set to 10 ppm and MS/MS tolerance was 0.8 Da for Orbitrap and linear ion trap, respectively. Carbamidomethylation of cysteine was set as a fixed modification. Oxidation of methionine and N-terminal Gln to pyro-Glu were set as variable modifications. We used a Mascot peptide significance threshold of *p* < 0.05 for post-search filtering. Quantitative and fold exchange values were calculated using Scaffold4 (Proteome Software, USA) for MS/MS-based proteomic studies.

### Quantitative reverse transcription (RT-q)PCR

RNA was extracted using ISOGEN II (Nippon Gene, Tokyo, Japan), and JEV RNA was quantified using a Power SYBR green RNA-to-Ct 1-Step kit (Thermo Fisher Scientific, Waltham, MA, USA) and the AriaMx Real-Time PCR system (Agilent, Santa Clara, CA, USA). The primers used were JEV forward 5′-AGCTGGGCCTTCTGGT’ and reverse 5′-CCCAAGCATCAGCACAAG’, and β-actin forward 5′-TTGCTGACAGGATGCAGAAG-3′ and reverse 5′-GTACTTGCGCTCAGGAGGAG-3′.

### Production of SRIPs

To generate JEV SRIPs, YFV-derived replicon plasmid (2.5μg, Yamanaka et al., 2021), PrME plasmid (1.25μg) and JEV core plasmid (1.25μg) were transfected into Huh7 WT or IPO7 KO cells. Each culture medium including SRIPs was harvested at 3 days post transfection. SRIPs for ZIKV, DENV and YFV were generated using replicon, PrME, core plasmid for each virus. Infectious titers were evaluated by the reporter activity on Vero cells infected with SRIPs.

### Statistical analysis

Normally distributed data (as determined using NORMDIST in Excel (Microsoft, version 16.75) were analyzed using two-tailed, unpaired parametric *t*-tests. Nonparametric data were analyzed using the Mann–Whitney U test or Kruskal–Wallis test followed by Dunn’s multiple comparison tests. Statistical analysis was performed using the GraphPad PRISM software (version 9.0.0). Significance was set to *p* < 0.05. Experimental schemes were prepared using BioRender.

## Acknowledgments

We would like to thank Ryoko Tanaka, Yuiko Azuma and Yuko Ookawa for secretarial work. We also thank Tomoko Maeda for helpful discussion. We thank the core instrumentation facility of the Research Institute for Infectious Diseases at Osaka University for MS analysis. This work was supported by the Japan Agency for Medical Research and Development (AMED), Research Program on Emerging and Re-emerging Infectious Diseases under JP20fk0108144, JP21fk0108144, and JP22fk0108144 (Y.Miyamoto and T.O.), JP23fk0108658 (T.O.), JP20im0210627, and JP22ym0126809 (Y.Miyamoto), 23fk0108656 (R.S) Life Science Foundation of Japan (Y.Miyamoto), Mochida Memorial Foundation for Medical and Pharmaceutical Research (T.O.), the Ministry of Education, Culture, Sports, Science, and Technology of Japan under 20H03495 (T.O.), and Boehringer Ingelheim research fund (T.O.).

## Author contributions

Y.Miyamoto. and T.Okamoto. designed the study; M.T., Y.Miyamoto. and Y.I. mainly performed experiments. A.T., M.M., and R.S. conducted the SRIP experiments. A.N., and F.S. conducted MS analysis. Y.Miyamoto., Y.I., and T.Okamoto wrote the manuscript. All authors discussed the results and commented on the manuscript.

